# NRF2 Dysregulation in Mice Leads to Inadequate Beta-Cell Mass Expansion during Pregnancy and Gestational Diabetes

**DOI:** 10.1101/2023.08.28.555207

**Authors:** Sharon Baumel-Alterzon, Isabelle Tse, Fatema Heidery, Adolfo Garcia-Ocaña, Donald K. Scott

## Abstract

The late stages of the mammalian pregnancy are accompanied with increased insulin resistance due to the increased glucose demand of the growing fetus. Therefore, as a compensatory response to maintain the maternal normal blood glucose levels, maternal beta-cell mass expands leading to increased insulin release. Defects in beta-cell adaptive expansion during pregnancy can lead to gestational diabetes mellitus (GDM). Although the exact mechanisms that promote GDM are poorly understood, GDM strongly associates with impaired beta-cell proliferation and with increased levels of reactive oxygen species (ROS). Here, we show that NRF2 levels are upregulated in mouse beta-cells at gestation day 15 (GD15) concomitant with increased beta-cell proliferation. Importantly, mice with tamoxifen-induced beta-cell-specific NRF2 deletion display inhibition of beta-cell proliferation, increased beta-cell oxidative stress and elevated levels of beta-cell death at GD15. This results in attenuated beta-cell mass expansion and disturbed glucose homeostasis towards the end of pregnancy. Collectively, these results highlight the importance of NRF2-oxidative stress regulation in beta-cell mass adaptation to pregnancy and suggest NRF2 as a potential therapeutic target for treating GDM.

## INTRODUCTION

The late stages of the mammalian pregnancy are accompanied with increased insulin resistance due to the increased glucose demand of the growing fetus. To maintain the maternal normal blood glucose levels, beta-cell mass expands leading to increased insulin release. Postpartum, beta-cell mass returns to its original size^1,2^. Beta-cell proliferation, beta-cell hypertrophy, beta-cell neogenesis, and protection from beta-cell apoptosis, are believed to drive human and rodent maternal beta-cell mass expansion, though the exact contribution of each mechanism is still in debate^3,4^. These processes are orchestrated by a specific combination of gestational hormones, growth factors and neurotransmitters. Of the most characterized are the lactogens (prolactin, placental lactogens), serotonin, hepatocyte growth factor (HGF), epidermal growth factor receptor (EGFR), estrogen, and progesterone. Once bound to their receptors, these factors trigger multiple mitogenic signaling pathways resulting in the expansion of beta-cell mass^1,3,5^.

Defects in beta-cell adaptive expansion during pregnancy can lead to gestational diabetes mellitus (GDM)^1^. With an estimation of 7.6% of the total pregnancy cases within the U.S, GDM may result in increased maternal risk for developing type 2 diabetes postpartum and in increased risk for fetal developmental disorders^6,7^. Various risk factors had been identified for GDM, including obesity, hyperlipidemia, hypertension, advanced maternal age, non-white race, family history of diabetes and polycystic ovary syndrome^6,7^. Importantly, although the exact mechanisms that promote GDM are poorly understood, GDM strongly associates with impaired beta-cell proliferation and with increased levels of reactive oxygen species (ROS)^1,8–11^.

Low levels of ROS are highly beneficial to beta-cells. Accordingly, they can improve beta-cell function and stimulate beta-cell proliferation^12–14^. However, when exposed to super-physiological ROS levels, beta-cells are presented with defects in insulin secretion, reduction in insulin content, decreased beta-cell identity, reduced beta-cell proliferation and increased beta-cell apoptosis, all of which result in the reduction of functional beta-cell mass^12,15^. Therefore, to protect themselves against oxidative stress potential damage upon exposure to high ROS levels, beta-cells activate the nuclear factor erythroid 2-related BZIP transcription factor 2, also known as NRF2 (gene name *Nfe2l2*), to maintain redox-balance^12^. Once activated, the NRF2 pathway can increase beta-cell survival, improve beta-cell function, maintain beta-cell identity, and promote beta-cell proliferation, to increase functional beta-cell mass^12,15^. In line with that, several SNPs in *NRF2* are linked to diabetes in GWAS studies ^16–19^.

Of note, while most studies on the physiological role of NRF2 in beta-cells were done in the context of Type 1 or Type 2 diabetic models, no study has uncovered the role of NRF2 in beta-cell expansion during pregnancy. We have previously shown that NRF2 is essential for beta-cell expansion and for maintenance of glucose homeostasis during insulin resistance following HFD feeding in mice^15^. Therefore, we hypothesized that NRF2 regulates beta-cell proliferation and survival during pregnancy, and thus contributes to maintain appropriate physiological beta-cell mass.

Here, we show that NRF2 levels are upregulated in mouse beta-cells at gestation day 15 (GD15) concomitant with increased beta-cell proliferation. Importantly, mice with tamoxifen-induced beta-cell-specific NRF2 deletion display inhibition of beta-cell proliferation, increased beta-cell oxidative stress and elevated levels of beta-cell death at GD15. These outcomes result in attenuated beta-cell mass expansion and disturbed glucose homeostasis towards the end of pregnancy. Taken together, NRF2 is an essential regulator of adaptive maternal beta-cell expansion during pregnancy and as a potential therapeutic target for treating GDM.

## RESULTS

### *Nrf2* expression is upregulated in mouse beta-cells at gestational day 15

Dual proteomics combined with RNAseq analysis have predicted that the transcription factor NFE2L2 (NRF2) could be an upstream regulator of gene expression changes observed in isolated mouse islets at GD14.5, which is a time when maximal beta-cell proliferation occurs in rodents^20^. However, these studies did not characterize the expression levels of NRF2 in islets during gestation nor did they study the role of NRF2 in beta-cell adaptation during pregnancy. Therefore, we first characterized NRF2 levels in beta-cells throughout pregnancy at gestational days GD11, GD15, GD19 or 4 days postpartum (PPD4) in C57BL6 mice by performing immunolabelling of NRF2 and insulin in pancreas sections from these mice. As shown in **Fig. 1a,b**, NRF2 levels significantly increased (10-fold) in beta-cells at GD15 compared to non-pregnant mice. However, NRF2 levels dramatically decreased during later stages of pregnancy (72% reduction in GD19 compared to GD15), returning to their basal levels postpartum (82% reduction in PPD4 compared to GD15). These results indicate that NRF2 is upregulated during pregnancy at a time when maximal beta-cell proliferation occurs and decreases over time when beta-cell mass returns to non-pregnant levels.

**Figure 1:**
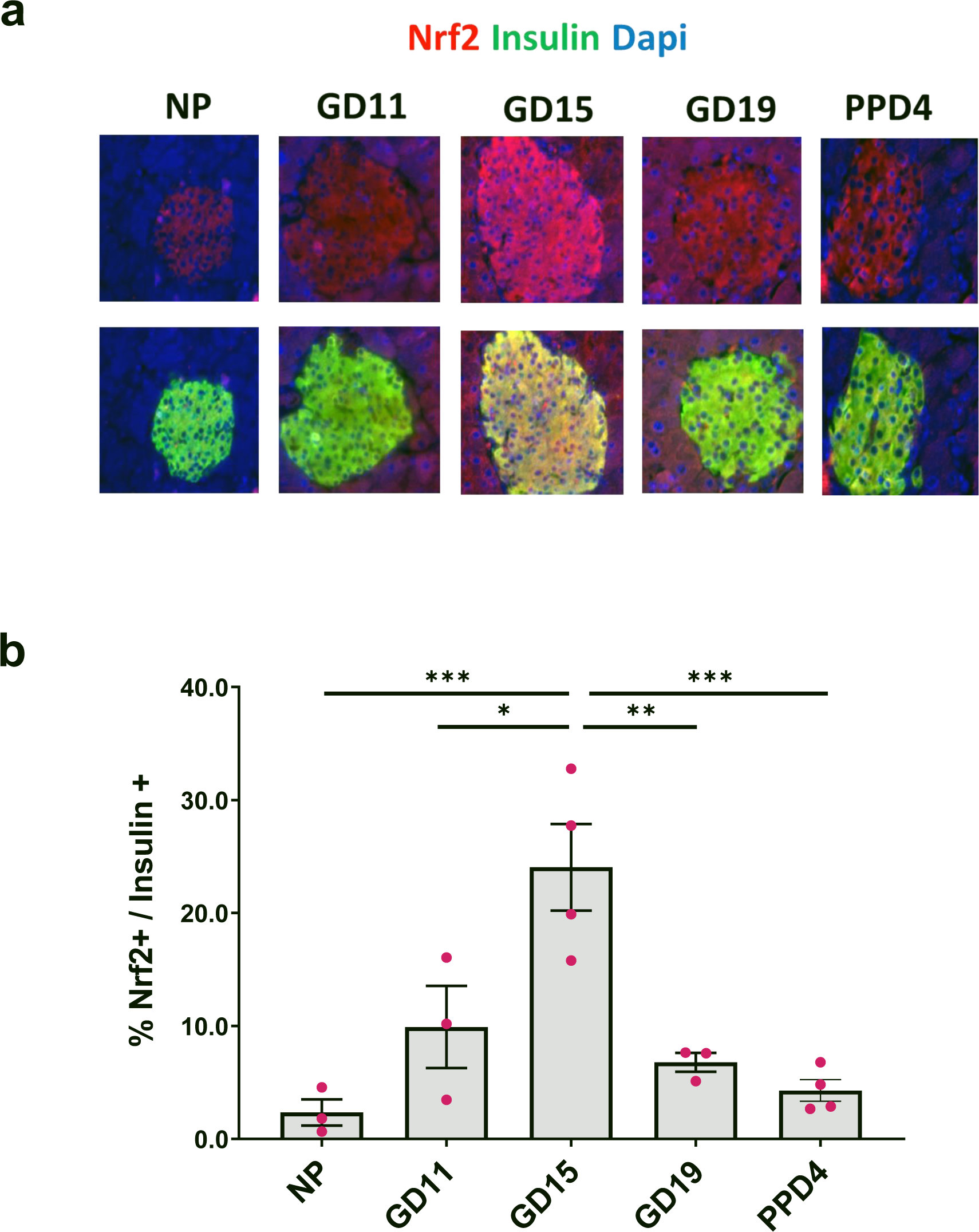
NRF2 levels are upregulated in mouse beta-cells at GD15. (**a,b**) Pancreatic sections from C57BL female mice at different gestational days were embedded and immunolabeled for NRF2 and insulin (scale bar 50 µM). Percentage of nuclear NRF2- and insulin-positive cells were calculated. Data are the means ± SEM for n=3-4, \**p* < 0.05, \*\**p* < 0.01, \*\*\**p* < 0.001.

### Beta-cell specific NRF2 deletion reduces beta-cell proliferation, increases beta-cell death and attenuates beta-cell redox balance at gestational day 15

As a key driving force in rodent maternal beta-cell expansion, beta-cell proliferation gradually increases during early stages of pregnancy, culminating at gestational days 13-15, followed by a progressive decline that reaches basal levels at gestational day 19^21,22^. Since NRF2 expression follows a similar pattern, this encouraged us to explore whether NRF2 plays a role in beta-cell proliferation during pregnancy. To investigate the link between NRF2 and beta-cell proliferation in pregnancy, we used our previously established Tam-induced beta-cell-specific NRF2 deletion mouse model (βNrf2KO)^15^. Due to the fact that administration of tamoxifen can lead to abortions and pregnancy complications^23^, female mice were treated for 5 consecutive days with tamoxifen [or corn oil vehicle (Veh) control] ip injections^15^ and then a 30 day period was established to allow tamoxifen to washout before initiating their overnight mating with males. Blood glucose, plasma insulin and intraperitoneal glucose tolerance test (IPGTT) were performed, and pancreata were harvested at the end of several gestational periods (GD15, GD19 or PPD4) (**Fig 2a**). Immunolabeling for NRF2 and insulin showed a 84% decrease in beta-cell NRF2 levels in pancreata harvested from GD15 Tam-injected βNrf2KO mice compared to Veh-injected mice (**Fig 2b,c**). In addition, there was no difference in the average number of embryos per mouse between Tam and Veh-injected mice (**Fig 2d**). Ki67 and insulin immunolabeling of Veh-injected mice showed the expected increase in beta-cell proliferation at GD15 compared to NP mice (9-fold), which then dropped to basal levels (**Fig 2e,f**). However, the increase of beta-cell proliferation at GD15 was blunted in Tam-injected βNrf2KO mice, indicating that NRF2 plays an important role in supporting beta-cell proliferation during pregnancy.

**Figure 2:**
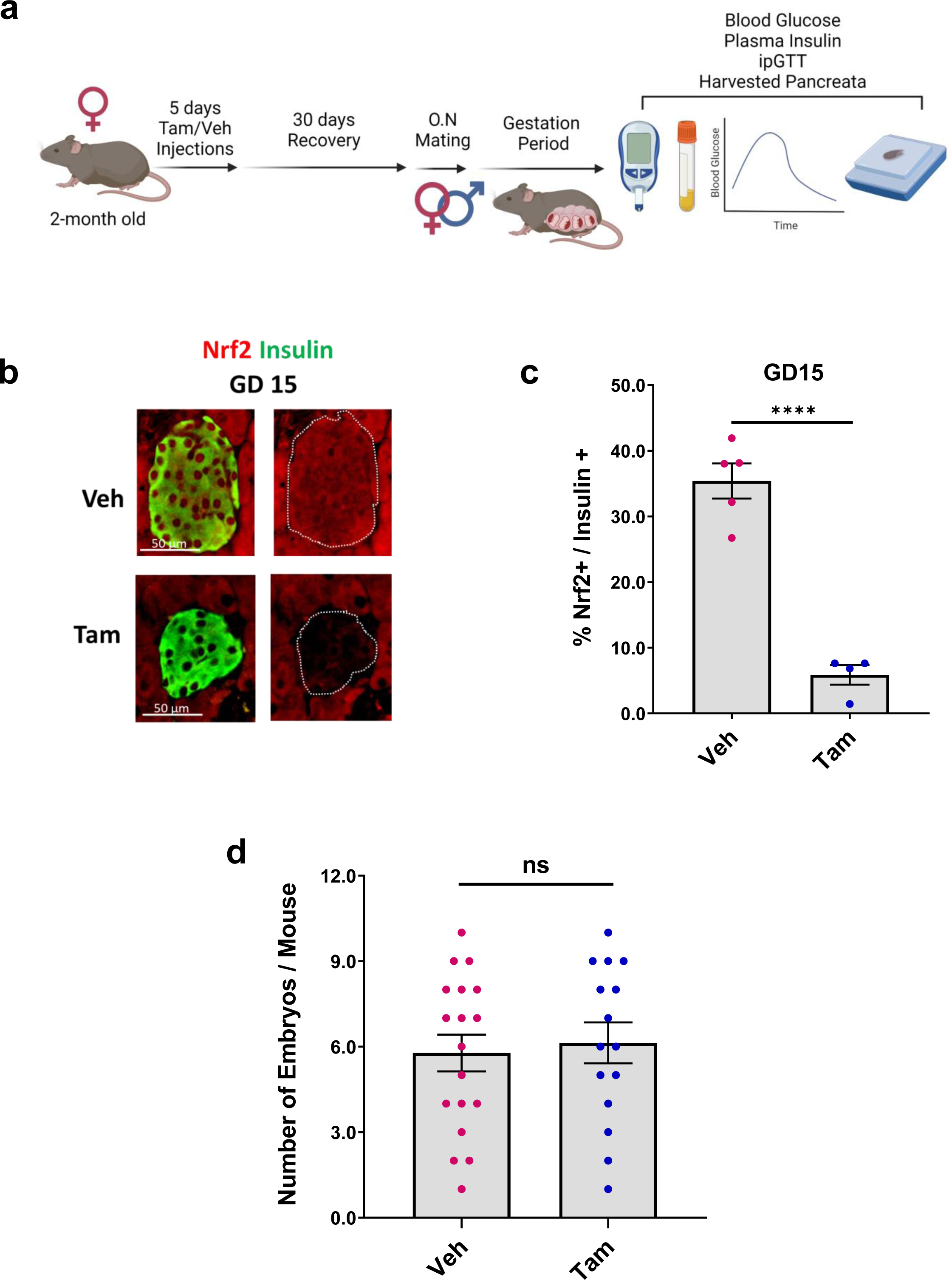
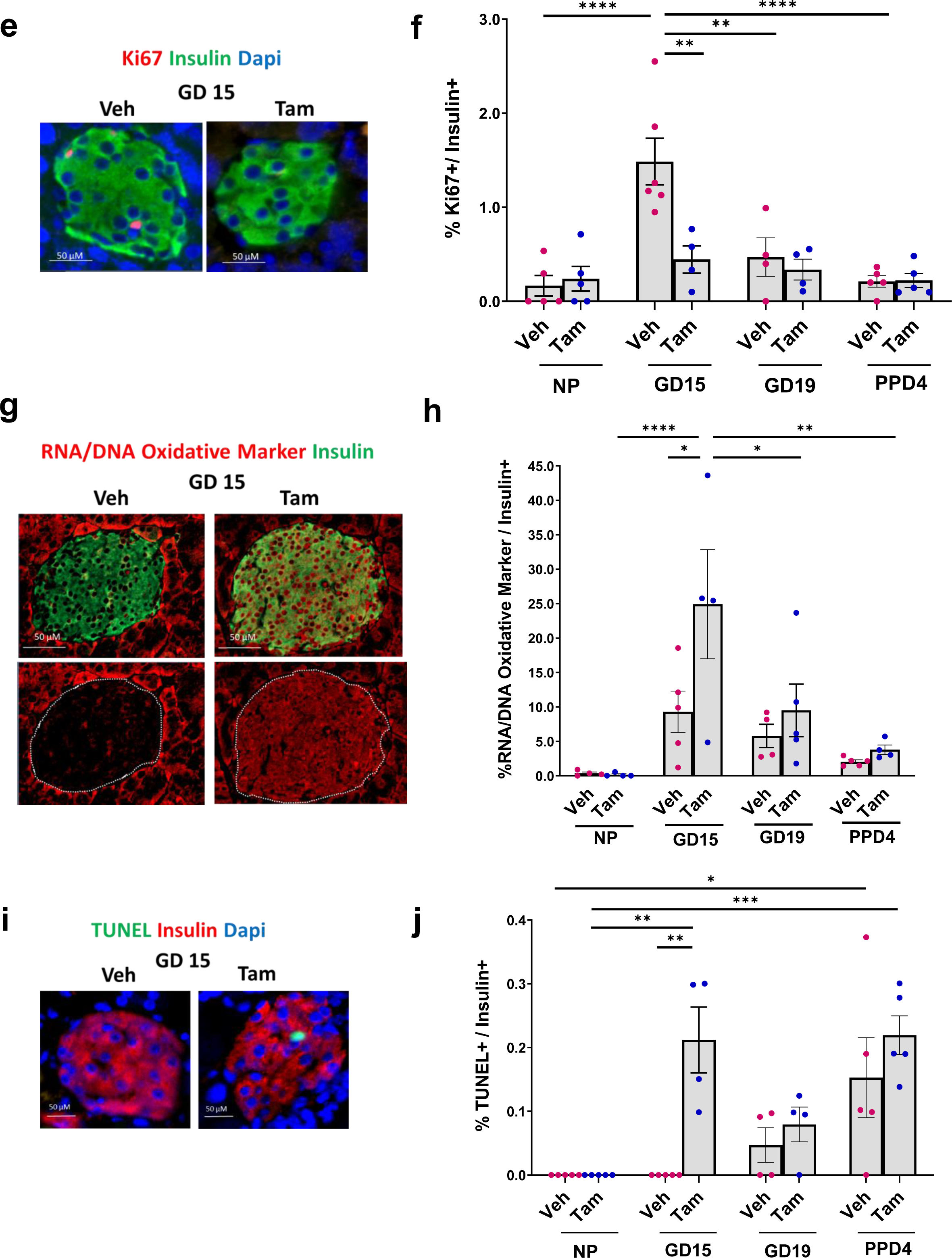
Beta-cell specific NRF2 deletion reduces beta-cell proliferation, decreases beta-cell survival and attenuates beta-cell redox balance at gestational day 15. (**a**) 2-month-old female beta-cell-specific NRF2 deletion mice were daily injected with tamoxifen (Tam) or corn oil (Veh) for 5 days. Following by 30 days of recovery period, females were placed overnight with males for mating. At the end of the gestational period (GD15, GD19 or PPD4), blood glucose and plasma insulin were taken, glucose tolerance test (ipGTT) was performed and pancreata were harvested for further immunolabelings (**b,c**). Pancreatic sections from GD15 Tam and Veh-injected females were immunolabeled for NRF2 and insulin (scale bar 50 µM). Percentage of nuclear NRF2- and insulin-positive cells were calculated. (**d**) Number of embryos per mouse was counted for Tam and Veh-injected pregnant females at different gestational days. (**e,f**) Pancreatic sections from Tam and Veh-injected females at different gestational days were immunolabeled for Ki67 and insulin (scale bar 50 µM), or (**g,h**) oxidative stress marker (RNA/DNA oxidative marker) and insulin (scale bar 50 µM), or (**i,j**) or TUNEL and insulin (scale bar 50 µM). Percentage of Ki67-positive or TUNEL-positive or oxidative stress marker-positive and insulin-positive cells were calculated. (**k,l**) Data are mean ± SEM for n=4-18. \**p* < 0.05, \*\**p* < 0.01, \*\*\**p* < 0.001, \*\*\*\**p* < 0.0001. (Fig 2a was created using BioRender.com).

Since NRF2 is a master regulator of antioxidant response in beta-cells^12^, we next tested whether deletion of NRF2 in beta-cells would result in changes of redox balance. Accordingly, we performed immunolabeling of pancreatic sections from these mice with antibodies recognizing either insulin or the oxidized nucleic acid species 8-OHdG, 8-OHG, 8-oxo-Gua, hence being used as a common RNA/DNA oxidative stress marker^15^ (**Fig 2g,h**). We found a small non-significant increase of oxidative stress in beta-cells of Veh-injected mice at GD15 compared to beta-cells of Veh-injected non-pregnant mice. Importantly, at GD15, Tam-injected βNrf2KO mice displayed a remarkable increase of oxidative stress in beta-cells (2.7-fold compared to GD15 in Veh control mice) that was reduced with time. These findings highlight NRF2’s role in maintaining adequate redox balance in beta-cells during pregnancy.

We then hypothesized that deletion of NRF2 in βNrf2KO mice would promote beta-cell death due to increased oxidative stress. To test this hypothesis, we immunolabeled pancreata for insulin and performed TUNEL assay (**Fig 2i,j**). In agreement with beta-cells undergoing a surge of beta-cell death postpartum^21,22,24^, beta-cells from both the Veh and Tam-injected βNrf2KO mice displayed increased beta-cell death at PPD4. Since ROS levels were downregulated postpartum, we speculate that this death was not a direct result of oxidative stress. However, at GD15, only Tam-injected βNrf2KO mice displayed increased beta-cell death (100% increase compared to Veh controls) that was associated with increased oxidative stress. This finding suggests that by regulating redox balance, NRF2 protects beta-cells from cell death at GD15.

### Beta-cell specific NRF2 deletion blunts beta-cell adaptive expansion and worsens glucose homeostasis towards the end of pregnancy

In order to test whether NRF2 plays a role in beta-cell mass expansion during pregnancy, pancreata from pregnant Veh and Tam-injected βNrf2KO mice were immunolabeled for insulin and analyzed for beta-cell mass by histomorphometry analysis (**Fig 3a,b**). Beta-cell mass reaches its maximal expansion in pregnant rodents at GD19^21,22^. Accordingly, Veh-injected mice displayed a significant increase in beta-cell mass compared to NP Veh mice (2.4 fold) at GD19. Yet, this increase was not observed in Tam-injected βNrf2KO mice, suggesting that NRF2 is essential for beta-cell mass adaptive expansion during pregnancy.

**Figure 3:**
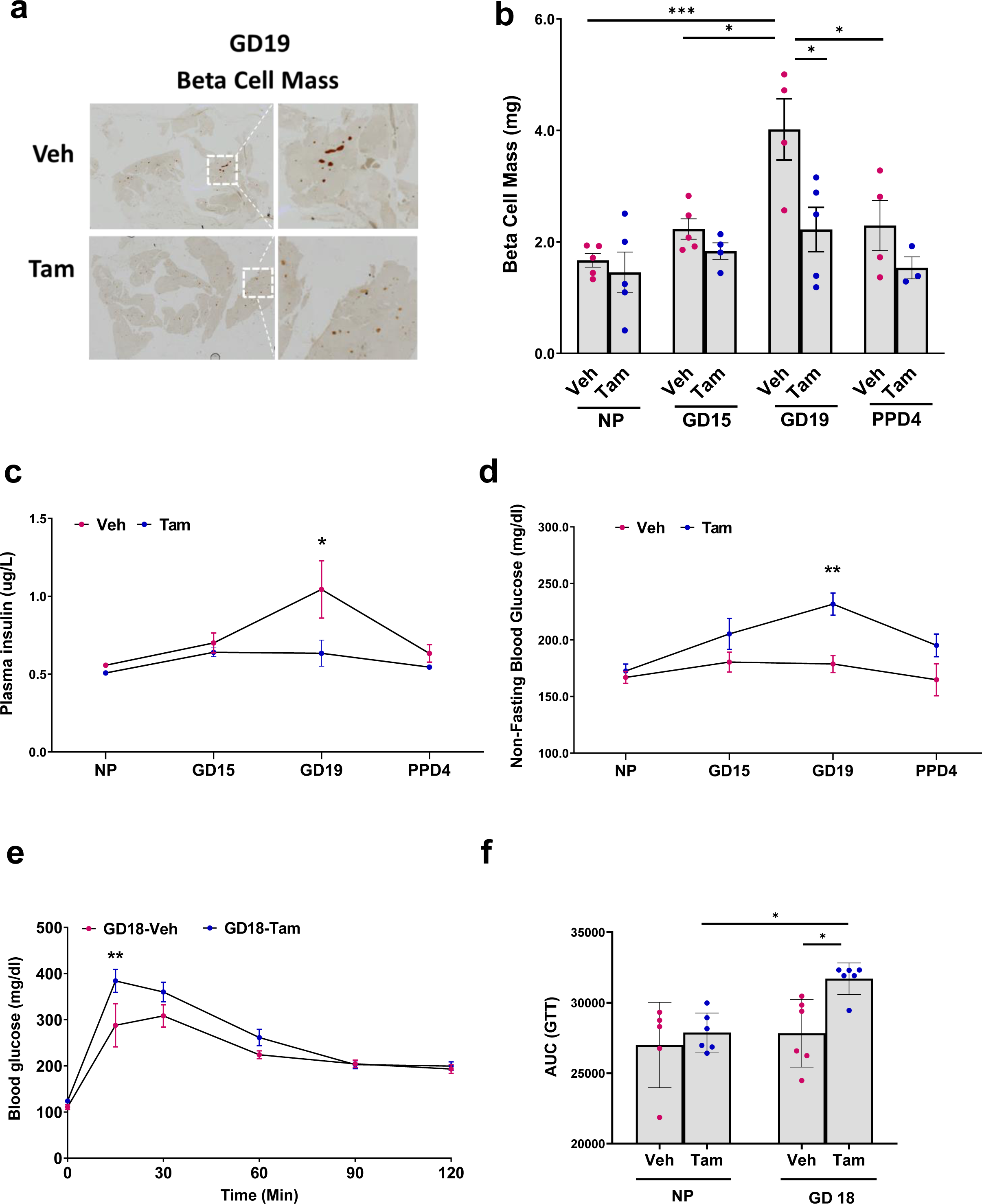
Beta-cell specific NRF2 deletion blunts beta-cell adaptive expansion and worsens glucose homeostasis towards the end of pregnancy. (**a,b**) Pancreatic sections from Tam and Veh-injected pregnant females at different gestational days were immunolabeled for insulin. beta-cell mass was calculated. (**c**) plasma insulin, (**d**) non-fasting plasma glucose were measured at different gestational days and (**e,f**) IPGTT was performed at GD18. Data are mean ± SEM for n=3-6. \**p* < 0.05, \*\**p* < 0.01, \*\*\**p* < 0.001.

To further explore the effect of beta-cell specific NRF2 deletion on glucose homeostasis, plasma insulin (**Fig 3c**), non-fasting blood glucose (**Fig 3d**) were collected at different gestational days. In addition, glucose tolerance test was performed at GD18 (**Fig 3e, f**). As expected, the increase in beta-cell mass in Veh-injected mice correlated with an increase in plasma insulin, improved glucose tolerance and sustained normal blood glucose levels. Conversely, inhibition of beta-cell mass expansion in Tam-injected βNrf2KO mice led to reduced plasma insulin levels, decreased glucose tolerance and hyperglycemia. These findings indicate that NRF2 is required for normal glucose homeostasis during pregnancy.

## DISCUSSION

Human pregnancy is accompanied by low levels of circulating ROS that are generated as a result of accelerated metabolism and due to heavy consumption of fatty acids that serve as a major energy source. At these low levels, ROS supports tissue remodeling and growth^11,25^. As in other tissues, low ROS levels are highly beneficial to beta-cells, improving insulin secretion, improving beta-cell identity, and even stimulating beta-cell proliferation^12^. In agreement with that, at GD15, maternal beta-cells of control pregnant mice display a small increase of ROS levels simultaneously with increased beta-cell proliferation. Thus, generation of low ROS levels during pregnancy likely contributes to the adaptive expansion of beta-cell mass. It is important to remember that since any increase in ROS levels inhibits KEAP1-dependent NRF2 degradation^12^, the small increase of ROS levels in maternal beta-cells at GD15 and potentially earlier, likely causes the increase in NRF2 levels during pregnancy. However, we cannot exclude the fact that growth factors and gestational hormones that are involved in the adaptive expansion of maternal beta-cells during pregnancy might affect NRF2 expression and activity. In fact, hepatocyte growth factor (HGF) and its receptor, c-Met, whose levels are increased at GD15, activate the NRF2 signaling pathway in human renal cancer cells and primary mouse hepatocytes^22,26,27^. Similarly, estrogen and its receptor, estrogen receptor-α (ERα), which play a role in beta-cell expansion during pregnancy; activate the NRF2 pathway in beta-cells^3,28^. Since prolactin, HGF, estrogen and serotonin have been shown to confer protection against oxidative stress^26,27,29–33^, it could be possible that they do it through regulation of NRF2 levels, an aspect that needs to be further studied.

Unlike low levels of ROS, high concentrations of ROS become toxic to beta-cells, inhibiting beta-cell proliferation, reducing beta-cell identity, hampering beta-cell function, and stimulating beta-cell apoptosis^12^. This highlights the importance of NRF2 as the master regulator of oxidative stress in pancreatic beta-cells. Not surprisingly, here we show that maternal beta-cells of pregnant βNrf2KO mice display high levels of ROS concomitant with reduced beta-cell proliferation and increased beta-cell death, all of which reduced beta-cell mass and worsened glucose homeostasis (**Fig 4**). We previously showed that NRF2 is necessary for beta-cell proliferation, survival, and expansion in male mice under conditions of obesity and metabolic stress^15^. While NRF2 plays a similar role in promoting beta-cell mass adaptive expansion in both obesity and pregnancy, more studies are needed to uncover the mechanism used by NRF2 to exert its action in these conditions. Of note, while beta-cell proliferation is the major driving force in maternal rodent beta-cell expansion during pregnancy, several indications suggest that beta-cell hypertrophy and beta-cell neogenesis might also contribute to this expansion^4,21,34^. However, whether NRF2 plays a role in these processes during pregnancy is unknown.

**Figure 4:**
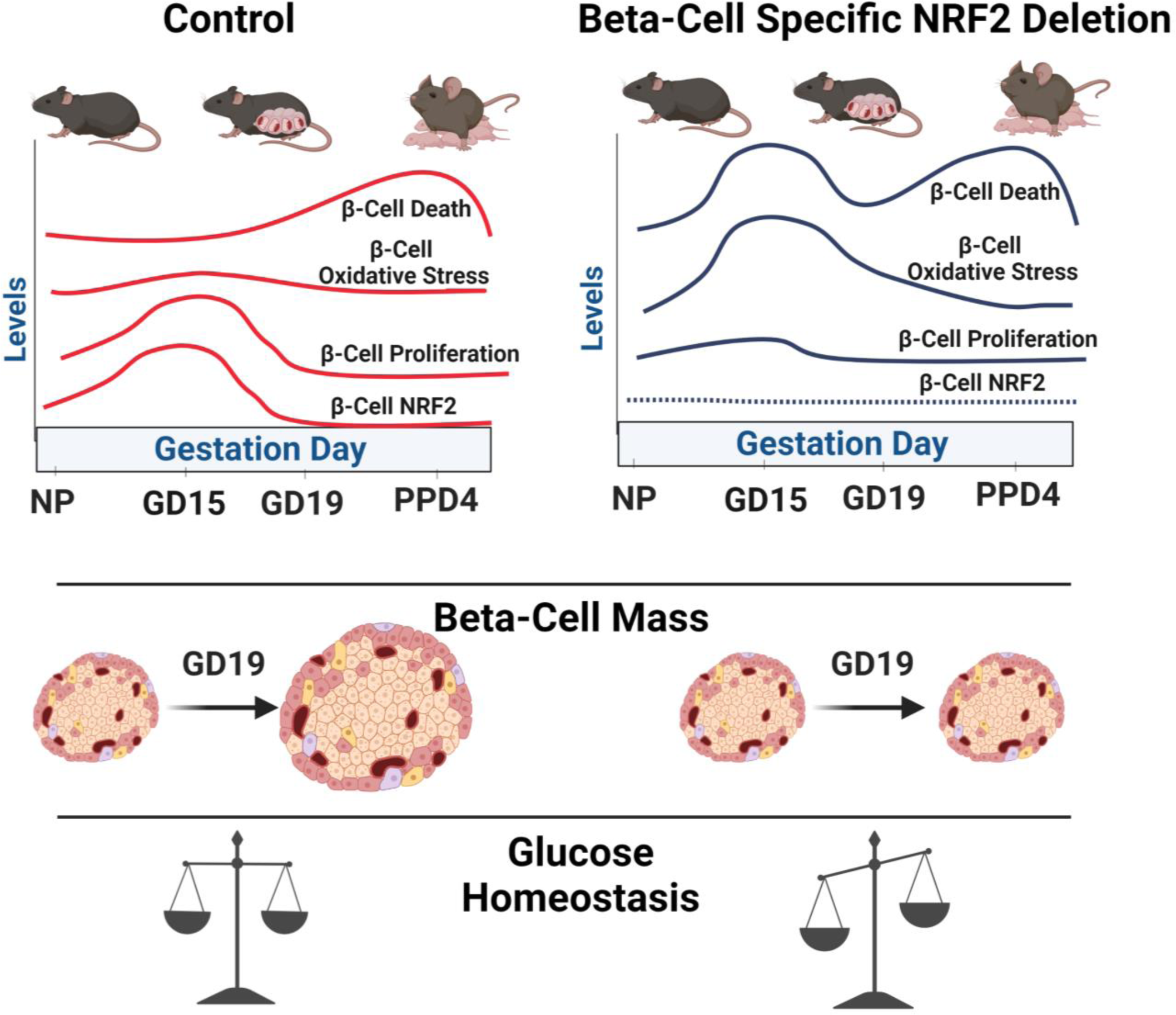
NRF2 stimulates beta-cell proliferation, beta-cell survival and beta-cell mass expansion during pregnancy. Beta-cell specific NRF2 deletion leads to high ROS levels, increased beta-cell death and reduced beta-cell proliferation at GD15. This results in attenuated beta-cell mass expansion at GD19 and disturbed glucose homeostasis. (Fig 4 was created using BioRender.com).

Multiple studies have linked GDM with dysregulation of the NRF2 signaling pathway^35^. However, while most of these studies focused on the placenta, umbilical cord, and adipose tissue, none has determined the role of NRF2 in beta-cell expansion and glucose homeostasis during pregnancy^35,36^. Our results show that beta-cell specific NRF2 deletion in pregnant mice leads to a GDM-like phenotype. This is likely due to reduced beta-cell proliferation, increased oxidative stress, accelerated beta-cell apoptosis and attenuated expansion of beta-cell mass. It could also be possible that beta-cell identity or function are directly affected by the deletion of NRF2 during pregnancy and studies to decipher their contribution to the observed effects are warranted.

Overall, the results of this study suggest that NRF2 can serve as a potential therapeutic target for treating GDM. Indeed, the NRF2 pharmacological activator, tertiary butylhydroquinone (THBQ) improves glucose homeostasis in a mouse model of GDM^35,36^. However, whether this is the result of alteration in the phenotype of beta-cells has not been studied. Notably, prevention of GDM using broad spectrum antioxidants is still controversial. Accordingly, while treatment with N-Acetyl-l-cysteine (NAC), vitamin E and vitamin C have been reported to ameliorate diabetes in GDM and type 2 diabetic mouse models^37–41^, clinical trials in women with GDM are still lacking and administration of NAC to type 2 diabetic patients did not improve their glycemia^42^. This could result from differences in the physiology between rodents and humans. Importantly, unlike the activation of endogenous NRF2 signaling, a pathway that is controlled by intrinsic feedback, administration of broad-spectrum antioxidants may eliminate even low level of ROS, which is actually beneficial to beta-cells. Future studies are needed to test the effect of NRF2 pharmacological activators on beta-cell mass expansion and glucose homeostasis in GDM pre-clinical mouse models.

## ACKNOWLEDGMENTS

We want to thank Dr Baumel-Alterzon’s K01 advisory committee members. Particularly, we would like to express our appreciation to Dr. Emily Bernstein from the Icahn School of Medicine at Mount Sinai and Dr. Sarah England from the Department of Obstetrics & Gynecology at Washington University School of Medicine for their valuable guidance and input on this study.

## DATA AVAILABILITY

The mouse lines generated in this study are available from the corresponding author upon reasonable request.

## AUTHORS’ RELATIONSHIPS AND ACTIVITIES

The authors declare that there are no relationships or activities that might bias, or be perceived to bias, their work.

## AUTHOR CONTRIBUTIONS

SBA, AGO, DKS developed the concept and designed the experiments. SBA analyzed the data. SBA, IT, FH conducted experiments and collected data. SBA drafted the manuscript. SBA, DKS, AGO, contributed to interpretation of data and critically revised the manuscript. DKS and AGO supervised the project. All authors gave their approval of the version to be published. SBA is the guarantor of this work and takes full responsibility for the content of the manuscript.

## METHODS

### Mouse Models

Females beta-cell specific NRF2 deletion mice were generated by crossing MIP-CreER^TAM^ mice^43^ (RRID: IMSR_JAX:024709) with Nrf2^lox/lox^ mice^44^ (RRID: IMSR_JAX:025433) as previously described^15^. Female mice were injected intraperitoneally for 5 consecutive days with 75 mg/g tamoxifen (Tam) (no. T5648; Sigma-Aldrich) dissolved in corn oil. Followed Tam injections, females were let recover for 30 days to allow Tam washout before mating. All studies were performed with the approval of and in accordance with guidelines established by the institutional animal care and use committee of the Icahn School of Medicine at Mount Sinai.

### Immunolabeling

Paraffin-embedded pancreatic sections were immunolabeled for Invitrogen #MA5-14520 Ki67 and GTX39371 Genetex insulin (for beta-cell proliferation) or Cayman Chemicals #10214 NRF2 and insulin (for NRF2 levels) or Abcam #ab62623 DNA/RNA oxidative damage and insulin (for oxidative stress) antibodies. The percentage of Ki67/NRF2/oxidative stress-positive/insulin-positive events was calculated from the total insulin-positive cells. For beta-cell mass an average of three insulin-stained mouse pancreatic section samples were measured using ImageJ software (National Institutes of Health).

### TUNEL Assay

TUNEL labeling was performed according to the manufacturer instructions, using the DeadEnd Fluorometric TUNEL System (G3250; Promega). Samples were then immunostained with insulin antibody (GTX39371; Genetex). The percentage of TUNEL-positive/insulin-positive events was calculated from the total insulin-positive cells.

### Glucose Homeostasis

Blood glucose was determined by glucometer and plasma insulin by ELISA (10-1249-01; Mercodia). For intraperitoneal glucose tolerance tests (IPGTT), mice were fasted for 16 h and then intraperitoneally injected with glucose at a dose of 2 g/kg D-glucose.

### Statistical Analysis

Data are presented as means ± SEM. The number of biologically independent replicates (n) for each experiment are indicated in the figure legends. Statistical analysis was performed using unpaired two-tailed t test, one-way ANOVA, and two-way ANOVA with Tukey multiple comparison test using GraphPad Prism (version 8.4.3). A p value < 0.05 was considered statistically significant.

